# In vitro inactivation of SARS-CoV-2 with 0.5% povidone iodine nasal spray (Nasodine) at clinically relevant concentrations and timeframes using tissue culture and PCR based assays

**DOI:** 10.1101/2021.01.31.426979

**Authors:** S.P. Tucker, S. Goodall, J. Julander, M. Mendenhall, P. Friedland, P.L. Molloy

## Abstract

**BACKGROUND:** There has been considerable speculation regarding the potential of PVP-I nasal disinfection as an adjunct to other countermeasures during the ongoing SARS-CoV-2 pandemic. Nasodine is a commercial formulation of 0.5% PVP-I that has been evaluated for safety and efficacy in human trials as a treatment for the common cold, including a Phase III trial (ANZCTR: ACTRN12619000764134). This study presents the first report of the *in vitro* efficacy of this formulation against SARS-CoV-2.

**METHODS:** We conducted *in vitro* experiments to determine if the PVP-I formulation inactivated SARS-CoV-2 using two independent assays and virus isolates, and incorporating both PCR-based detection and cell culture methods to assess residual virus after exposure to the formulation.

**RESULTS:** Based on cell culture results, the PVP-I formulation was found to rapidly inactivate SARS-CoV-2 isolates *in vitro* in short timeframes (15 seconds to 15 minutes) consistent with the minimum and maximum potential residence time in the nose. The Nasodine formula was found to be more effective than 0.5% PVP-I in saline. Importantly, it was found that the formulation inactivated culturable virus but had no effect on PCR-detectable viral RNA.

**CONCLUSIONS:** The PVP-I formulation eliminated the viability of SARS-CoV-2 virus with short exposure times consistent with nasal use. PCR alone may not be adequate for viral quantification in nasal PVP-I studies; future studies should incorporate cell culture to assess viral viability. Nasal disinfection with PVP-I may be a useful intervention for newly-diagnosed COVID-19 patients to reduce transmission risk and disease progression to the lower respiratory tract.

## Introduction

SARS-CoV-2, the virus responsible for COVID-19 is reported to replicate effectively in the upper respiratory tract (URT) with an apparent preference for the nasopharynx which exhibits a higher viral load and a higher viral replication rate than the oral cavity and lower respiratory tract (LRT) ^1^ ^2^.Entry is mediated by two proteins, the angiotensin converting enzyme-2 (ACE2) receptor and the transmembrane protease serine 2 (TMPRSS2), which facilitate viral attachment and membrane fusion prior to entry into the cell ^3^. Consistent with the observed preferential URT replication, these proteins are abundant in nasal goblet secretory cells and nasal ciliated epithelium, and are also expressed in pulmonary type II pneumocytes in alveoli and ileal absorptive enterocytes ^4^.

Virus is readily isolated from the nose early in the course of the disease (in many cases prior to significant symptoms) and shedding from the nose/URT is thought to be a primary mode of transmission ^3^. Preferential infection via the nasal mucosa and subsequent dissemination via the URT suggest that the nose and associated URT are central to the entry of virus and subsequent spread, including spread to other tissues, such as the LRT, as well as new hosts ^3^. These observations suggest that the nasal cavity presents a target for early intervention with the goal of not only preventing and/or limiting transmission to others, but also reducing the risk of progression of the infection to the LRT, which could be considered a secondary stage of the disease.

Povidone-iodine or PVP-I is a complex of polyvinylpyrrolidone and iodine. PVP-I is considered to have a broader spectrum of antimicrobial action compared with other common antiseptics ^5^ ^6^ ^7^. PVP-I is routinely used as a general disinfectant of skin and mucous membranes and there have been numerous clinical studies demonstrating the safety of PVP-I in a variety of topical applications in ophthalmology, otology, rhinology and dermatology ^8^. The use of PVP-I in the nasal passages has been explored for decolonization of potentially pathogenic bacteria (PPB) and most recently has been proposed as an intervention to assist in the management of SARS-CoV-2 infection ^9^ ^10^. Pelletier and colleagues disclosed the *in vitro* inactivation of SARS-CoV-2 after a 60 second exposure to a series of proposed aqueous nasal and oral rinse PVP-I solutions ^11^. Others have proposed that PVP-I represents an excellent option for localized disinfection of the URT as a means of augmenting other precautionary practices during the COVID-19 pandemic and thereafter ^12^ ^13^ ^14^ ^15^ ^16^. However, none of the intranasal PVP-I formulations has been proposed has been developed for nasal use, rigorously tested for safety in human clinical trials or shown to be stable for commercially-meaningful periods at the concentrations needed for safe intranasal use.

The antimicrobial activity of PVP-I is dependent on the concentration of free iodine and the contact time with microorganisms ^17^. However, iodine is a reactive species that is relatively unstable in solution at low concentrations and incompatible with several plastic storage vessels. Disinfection of the nasal epithelium is complicated by rapid clearance of materials from the nasal cavity, due to nasal discharges and mucociliary clearance. Furthermore, high concentrations of PVP-I are unacceptable for nasal use due to reported ciliotoxicity, local sensitivity and the risk of iodine uptake through the nasal mucosa ^18^ ^19^ ^20^. The challenges for development of an effective intranasal formulation of a PVP-I nasal disinfectant are therefore the identification and administration of PVP-I in a formulation that is readily administered by users, stable for convenient distribution and use, effective as an antimicrobial despite very short exposure times, and safe for intranasal use.

Nasodine^®^ Antiseptic Nasal Spray (Nasodine) is a stable formulation of 0.5% PVP-I optimized for the safe and effective disinfection of the nasal epithelium. Preclinically, it has been evaluated for intranasal safety using a sensitive air-liquid interface (HNEC-ALI) model of human nasal epithelium^21^. Even with 30 minutes exposure time, Nasodine produced no ciliotoxicity, had no detrimental effects on the paracellular permeability and showed no indication of cellular toxicity. The authors concluded: “these results indicate a good safety profile of 0.5% povidone-iodine when applied to HNEC-ALI cultures *in vitro*”^21^.

Subsequently, the product was developed by the sponsor^a^ as a treatment for viral upper respiratory infections, with a focus on the common cold, and tested in three human studies for safety and efficacy, including a Phase III trial in adults with cold symptoms (ANZCTR: ACTRN12619000764134). The sponsor has reportedly filed a registration dossier with the Therapeutic Good Administration in Australia seeking its approval as an over-the-counter medicine for treatment of the common cold.

Early in the development of the product, its activity was confirmed *in vitro* against representative strains of all major viruses responsible for respiratory infections, including rhinovirus (hRV-14 and hRV-2), coronavirus (E229), parainfluenza virus, influenza A virus, respiratory syncytial virus (RSV) and metapneumovirus (unpublished data). In this report, we present the first data evidencing the *in vitro* efficacy of Nasodine against SARS-CoV-2.

## Materials and Methods

### CELLS AND VIRUS

Vero cells (ATCC^®^, CRL-1587, ATCC, Manassas, Virginia) were grown in Eagle’s Minimum Essential Medium (EMEM, Sigma-Aldrich, North Ryde, Australia, or Sigma-Aldrich, St. Louis, Missouri), supplemented with 1x non-essential amino acids (Gibco, Mount Waverley, Australia, or Thermo Fisher Scientific, 168 Third Avenue, Waltham, MA) and 10% heat-inactivated fetal bovine serum (FBS: Bovogen, Melbourne, Australia, or GE Healthcare Hyclone, Marlborough, MA). Vero E6 cells (ATCC^®^, C1008) were grown in MEM/EBSS with L-Glutamine (Hyclone) supplemented with 1X non-essential amino acids (Gibco), 1 mM sodium pyruvate (Sigma-Aldrich), and 5% characterized fetal bovine serum (Hyclone).

Two isolates of SARS-CoV-2 were used: 1) BetaCoV/Australia/VIC01/2020 was isolated and grown by the Victorian Infectious Diseases Reference Laboratory from a positive patient specimen in January 2020. Whole-genome sequencing confirmed the presence of SARS-CoV-2 (GenBank ID: MT007544). 2) SARS-CoV-2/USA_WA1/2020 was prepared by Natalie Thornburg, CDC and provided by WRCEVA, University of Texas Medical Branch. Virus stocks were prepared by growing virus in Vero 76 cells (ATCC CRL-1587) using media supplemented with 2% fetal bovine serum.

### POVIDONE IODINE SOLUTIONS

Nasodine Antiseptic Nasal Spray (0.5% povidone-iodine) was supplied by Firebrick Pharma Limited, Melbourne Australia.

### INCUBATION OF VIRUS WITH TREATMENT SOLUTIONS

Protocols 1 and 2 were as follows:

1. SARS-CoV-2 (BetaCoV/Australia/VIC01/2020 at 1.2 × 10^5^ TCID_50_ units/mL) was incubated in EMEM alone (negative control) or Nasodine at 37°C for 1 minute.
2. SARS-CoV-2 (USA_WA1/2020 at 2 × 10^5^ TCID_50_ units/mL) was incubated in saline (negative control), 0.5% PVP-I in saline, or Nasodine, each for 15 seconds, 5 minutes or 15 minutes. Reactions were quenched by 10-fold dilution using ice cold medium containing 2% or 10% FBS, respectively. Immediately after adding ice cold media, viral titers were determined via TCID_50_ assay and/or RNA copies estimated by RT-PCR.

### TCID50 ASSAY

Vero E6 cells were seeded into 96-well plates at a cell density of 2×10^4^ cells/well and incubated overnight at 37°C, 5% CO_2_ to 80-100% confluency, prior to infection with replicate serial dilutions of the reaction mix. Plates were incubated at 37°C, 5% CO2 atmosphere for 96 hours before harvesting and assay for viral RNA by RT-PCR (Protocol 1), or for 6 days prior to scoring for presence or absence of viral cytopathic effect (Protocol 2). Titers were calculated using the method of Reed and Meunch.

### PURIFICATION OF SARS-COV-2 RNA

SARS-CoV-2 RNA was purified using the QIAamp 96 virus QIAcube HT Kit and processed on the QIAcube robotic extraction platform (QIAgen, Hilden, Germany)

### TAQMAN RT-PCR OF SARS-COV RNA

Purified SARS-CoV-2 RNA was reverse transcribed using the Bioline Sensifast cDNA kit (Catalogue number CSA-01148; London, UK). Real-time assays were performed using the method of Corman and colleagues ^22^ with E gene primers and probes (IDT, Singapore) and ABI TaqMan Fast Universal PCR Master Mix (2x) (catalogue number 4352042; Thermofisher, Vilnius, Lithuania) on a Thermofisher ABI 7500 Fast Real Time PCR machine.

## Results

Two related protocols are described, the first associated with a preliminary study (Protocol 1) and the second (Protocol 2) was a confirmatory study in an independent laboratory, utilizing a different source of SARS-CoV-2.

Since cytopathic effects (CPE) associated with BetaCoV/Australia/VIC01/2020 were uncertain at the time of study, to confirm the presence or absence of replicating SARS-CoV-2, samples from all TCID50 plate wells were analyzed for the presence of SARS-CoV-2 RNA by Taqman RT-PCR. Confirmatory experiments under Protocol 2 involving SARS-CoV-2/USA_WA1/2020 used a CPE-based assay.

### Protocol 1

To establish a baseline of RT-PCR cycle threshold (Ct) values corresponding to input RNA copies prior to replication, TCID50 plates were prepared and assayed immediately without incubation (corresponding to the viral inoculum reaction mix at 0 h post infection). SARS-CoV-2 specific Ct values for inocula prepared from virus exposed to media alone or Nasodine (in duplicate) prior to replication on Vero cells showed a linear relationship between Ct-value and dilution factor (Figure 1A). The Ct values for Nasodine or control treated virus were indistinguishable at all dilutions indicating that Nasodine treatment did not interfere with the ability to detect nucleic acid by real-time RT-PCR and had no effect on the apparent number of RNA copies detected in the inocula.

**FIGURE 1.**
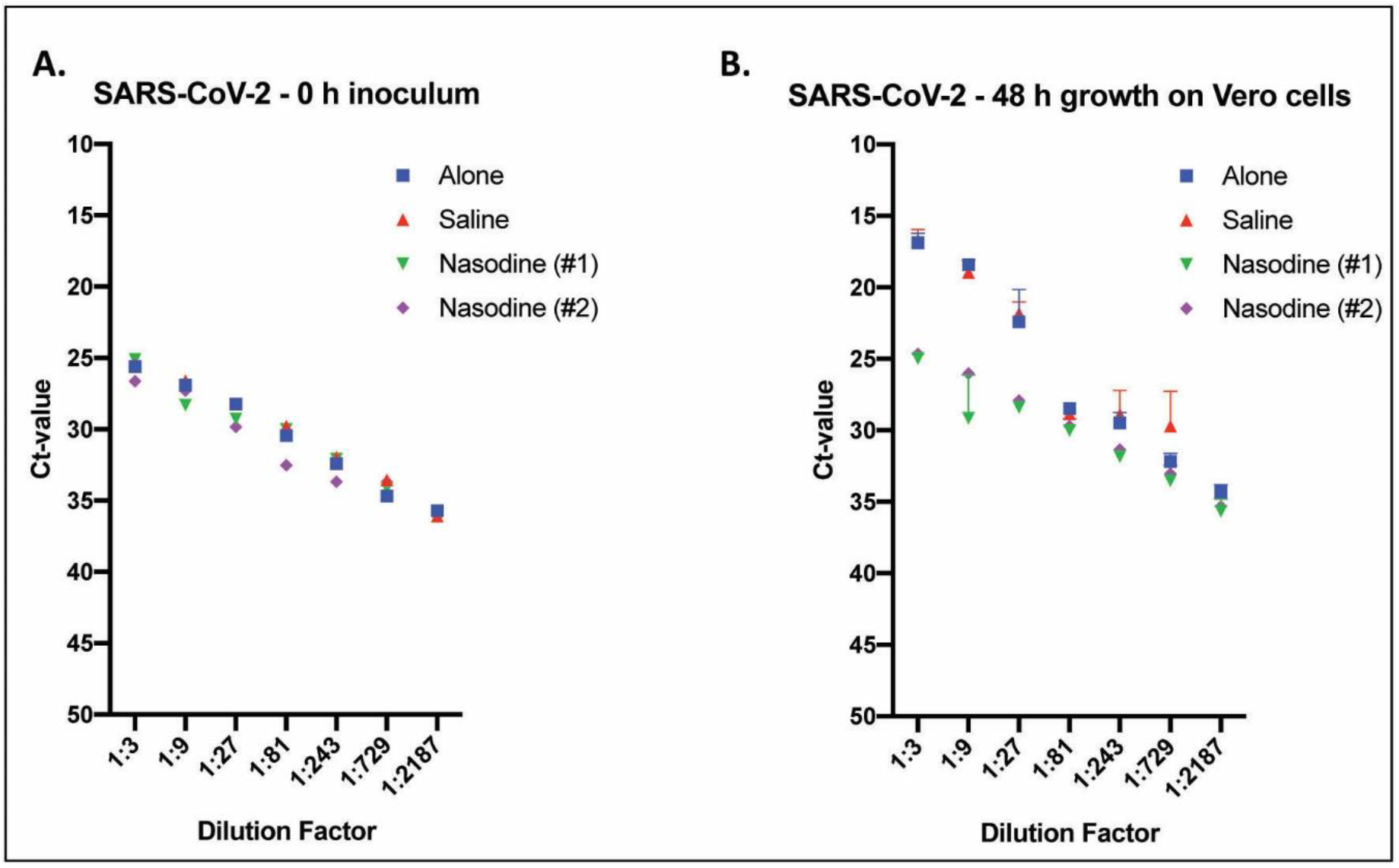
Titre of Nasodine and control treated SARS-CoV-2 via TCID_50_ assay and RNA detection via real-time TaqMan RT-PCR. SARS-CoV-2 was exposed to the indicated test solution(s) for 1 minute before serial dilution (1:3) and incubation on Vero cells for either 0 (zero) or 96 h. Values expressed as mean cycle threshold (Ct) value + SEM (n=3) versus dilution factor. (A) Time point zero (0 h) inoculum titration used to determine baseline Ct-values of treated samples prior to replication in Vero cells. (B) Titres associated with cultures harvested 96 h post inoculation of Vero cells.

SARS-CoV-2 treated with control media alone displayed robust replication in the TCID_50_ plates incubated for 96 h (Figure 1B), as measured by decreased Ct-values corresponding to increased RNA copies relative to the copies present in the inoculum. In contrast, the samples incubated in the presence of Nasodine displayed an apparently linear relationship between viral RNA Ct-values and dilution factor at 96 hours post-infection (Figure 1B). The Ct values associated with the Nasodine treated samples titred by TCID_50_ assay at 96 hours post-infection overlapped with the values obtained from the 0-hour inoculum assay indicating no apparent increase in RNA copies over 96 hours and, therefore no evidence of viral replication (Figure 1).

### Protocol 2

This protocol confirmed and extended the preliminary study data by using a different virus isolate (USA_WA1/2020), a CPE-based assay and three different incubation times: 15 seconds, 5 minutes and 15 minutes. Only the 15 second and 5 minute results were considered relevant to nasal use because of nasal clearance, due to mucous and mucociliary clearance, and likely inactivation of iodine by mucins and proteins in the nose.

Nasodine treatment resulted in a 3.5 log unit reduction in virus titer in 15 seconds, a 4.0 log unit reduction following 5 minute exposure and no detectable viable virus after 15 minute incubation (Table 1). Interestingly, Nasodine was found to be more effective than 0.5% PVP-I alone (prepared in saline) at both the clinically-relevant 15 second and 5 minute time-points.

**TABLE 1.** Virucidal efficacy against SARS-CoV-2 after incubation with virus at 37°C. Untreated control titers were 5.0, 5.0 and 5.3 log TCID_50_ units/mL at 0.25, 5 and 15 minutes, respectively. 70% ethanol was used as a positive control and reduced virus titer to baseline (0.67 log_10_ TCID_50_/ml).

| Treatment | Incubation time (minutes) | Virus titer (log<sub>10</sub> TCID<sub>50</sub> units/mL) | Change in titer compared to control (log<sub>10</sub> TCID<sub>50</sub> units/mL) |
| --- | --- | --- | --- |
| Nasodine | 0.25 | 1.5 | -3.5 |
| PVP-I saline | 0.25 | 2.5 | -2.5 |
| Saline | 0.25 | 5.0 | 0 |
| Nasodine | 5 | 1.0 | -4.0 |
| PVP-I saline | 5 | 1.67 | -3.3 |
| Saline | 5 | 4.5 | -0.5 |
| Nasodine | 15 | <0.67 | >-4.3 |
| PVP-I saline | 15 | <0.67 | >-4.3 |
| Saline | 15 | 5.5 | 0.2 |

## Discussion

Nasodine is a PVP-I based antiseptic nasal spray formulated for stability, convenience and tolerability for use in the nasal cavity. It has undergone all phases of clinical development inluding a Phase III study to assess its safety and efficacy as a treatment for the common cold (ANZCTR: ACTRN12619000764134). The advent of the COVID-19 pandemic resulted in speculation from several sources about the potential utility of PVP-I based antiseptics as an adjunct to existing personal protective equipment (PPE) together with numerous reports of *in vitro* activity and the initiation of clinical trials involving various formulations of PVP-I (see for example ^12^ and clinicaltrials.gov NCT04347954). We therefore set out to determine if the Nasodine formulation inactivated SARS-CoV-2 *in vitro* as a prelude to clinical studies in COVID-19 patients.

The data described herein demonstrate that Nasodine treatment results in the rapid inactivation of SARS-CoV-2 in short timeframes that are consistent with clinical utility. Preliminary experiments using BetaCoV/Australia/VIC01/2020 and a PCR based TCID_50_ assay found that 1 minute exposure resulted in the elimination of all detectable, replication-competent viable virus, indicating that Nasodine effectively inactivated 1.2 × 10^5^ TCID_50_ units/mL (5 log units) of SARS-CoV-2 with a 1-minute exposure. These data were confirmed using a different isolate (SARS-CoV-2/USA_WA1/2020) and an alternative CPE-based assay which showed 3.5 and 4.0 log reductions at 15 seconds and 5 minutes exposure, respectively.

In Protocol 1 of this study, Nasodine treatment had no effect on the detection of RNA or the estimated number of RNA copies detected in the samples prior to culture in Vero cells (0 hours) and in the treated samples obtained from Vero cell cultures at 96 hours post infection. These data suggest that amplifiable RNA remains after PVP-I treatment and that RT-PCR alone may not be an acceptable method for measuring the immediate effect of PVP-I on viral viability. It is possible that such residual RNA may be cleared by other mechanisms *in vivo*. These observations are consistent with the clinical findings of Lamas and colleagues who reported that PVP-I based mouthwash reduced SARS-CoV-2 RNA copies in COVID-19 patients’ saliva at 1-3 hours post treatment but not at the earlier time point of 5 minutes post treatment ^23^. This is an important finding in the context of several studies currently underway to assess the effectiveness of various PVP-I nasal formulations in the COVID-19 setting, which may be using PCR only as the measure of residual virus (see for example clinicaltrials.gov NCT04347954).

The virucidal effect of Nasodine on SARS-CoV-2 is consistent with findings reported by others using various other formulations of PVP-I on SARS-CoV-2 other coronaviruses *in vitro* (see for example ^9,24,25^). Nasodine thus offers the prospect of a broad-spectrum antiseptic/disinfectant for the nasal tract and is currently undergoing review by regulatory authorities in anticipation of use in the treatment of the common cold. In addition to the common cold, it may prove to have utility in the eradication of SARs-CoV-2 from the nasal passages as an adjunct to PPE in the management of COVID-19. We have subsequently started a first-in-human clinical trial to determine if these SARS-CoV-2 *in vitro* data translate to reduced viral shedding in COVID-19 patients. If successful, we anticipate that nasal disinfection with PVP-I may become a useful addition to the repertoire of PPE and other protections for health care workers; it also may present a useful intervention for newly-diagnosed COVID-19 patients, both to reduce transmission risk to others and to reduce the incidence of progression of the disease to the lower respiratory tract.

## Acknowledgements

Supported by Firebrick Pharma Ltd, Melbourne Australia

The authors thank Drs. Leon Caly and Julian Druce of the Virus Identification Laboratory, Victorian Infectious Diseases Reference Laboratory, 792 Elizabeth Street, Melbourne, Victoria, Australia, 3000 for kindly providing BetaCoV/Australia/VIC01/2020 and for conducting the assays described under Protocol 1.

## Footnotes

a Nasodine^®^ Antiseptic Nasal Spray is sponsored by Firebrick Pharma Ltd, Melbourne Australia.

